# Deep soil water as a dominant source for spruce water uptake in a subalpine forest: evidence from multi-year isotope data

**DOI:** 10.64898/2026.03.10.710581

**Authors:** Harsh Beria, Ankit Shekhar, Nina Buchmann, Mana Gharun

## Abstract

– Norway spruce (*Picea abies*) dominates many European mountain forests, yet their seasonal water uptake strategies in high-elevation mono-specific natural stands remain poorly understood. We quantified contributions of shallow (0-10 cm) and deep (50-70 cm) soil layers to tree water uptake over three consecutive growing seasons (2020-2022) using stable water isotopes and Bayesian mixing analysis.
– Contrary to the prevailing view of spruce as a shallow-rooted species relying primarily on water from the upper 10-20 cm of soil, our results showed more than 50% water uptake from deeper soil (50-70 cm), with deeper soil contributions crossing 80% in 2020.
– During the dry and warm summer of 2022, positive soil recharge and elevated atmospheric demand increased evapotranspiration, with spruce trees taking up recently infiltrated rainfall from different soil depths, including >50% uptake from deeper layers.
– Spruce water uptake shifted from cold-season-recharged soil water early in the growing season to warm-season precipitation in late summer. The timing of this shift in mid-summer can be explained by soil water recharge from recent rainfall infiltrated into the entire soil profile. This reliance on summer precipitation increases vulnerability of mono-specific spruce stands to more frequent droughts and heat waves under future climate change.

## Introduction

Mountain forests play an important role in regulating water and carbon dioxide fluxes, at the same time they remain highly sensitive to climatic variability. In these environments, ecohydrological processes are tightly linked to the timing and magnitude of liquid water inputs, via rainfall or snowmelt, which control soil water storage and, together with energy availability, regulate evapotranspiration. Understanding tree water uptake dynamics in such landscapes is essential for accurately predicting their responses to climatic change, especially because mountain regions are experiencing rapid warming (Pepin *et al*., 2015, 2022) and pronounced declines in snow cover (Matiu *et al*., 2021).

Subalpine forests of the European Alps are undergoing substantial greening (Filippa *et al*., 2019; Obuchowicz *et al*., 2023), which may enhance carbon uptake, while enhancing evapotranspiration (Mastrotheodoros *et al*., 2020). However, most ecohydrological studies investigating tree water uptake dynamics have focused on low-elevation subalpine forests (Bertrand *et al*., 2014; Brinkmann *et al*., 2018; Goldsmith *et al*., 2019; Floriancic *et al*., 2024; Bernhard *et al*., 2025; Zuecco *et al*., 2026), with a limited number of studies at higher elevations (Brighenti *et al*., 2024). This creates a substantial gap in our understanding of water-use strategies in high elevation subalpine forests, where increasing energy availability, earlier snowmelt, longer growing seasons, and complex topography are expected to alter ecohydrological processes in ways that differ fundamentally from lower-elevation forests.

Tree water uptake dynamics are commonly inferred using mixing models applied to stable isotopes of water in soil and xylem waters (Ehleringer & Dawson, 1992; Popp *et al*., 2025), to quantify the seasonal origin of plant water (e.g., winter vs. summer precipitation) and the depths from which trees extract water (e.g., shallow vs. deep soil). In the European Alps, coniferous trees, typically *Picea abies* (Norway spruce or “spruce” used hereafter) and *Larix decidua* (referred to as “larch” hereafter) are the prominent species. Larch, being a pioneering species, is usually present closer to the treeline, whereas spruce spans a wider elevation range. Many conifer species use winter precipitation or snowmelt (Martin *et al*., 2018; Langs *et al*., 2020; Zhang *et al*., 2021; Nehemy *et al*., 2022; Brighenti *et al*., 2024) present within the entire soil profile at the onset of the growing season and may either switch to summer precipitation and thus shallow soil depths (Martin *et al*., 2018; Berkelhammer *et al*., 2020; Sprenger *et al*., 2022; Brighenti *et al*., 2024) or continue using winter-recharged deep soil depths for water uptake (Zhang *et al*., 2021). These contrasting observations reflect differences in rooting depth, interspecies competition, soil type, and climatic aridity (Xiangyang *et al*., 2022; Sun *et al*., 2024; Hackmann *et al*., 2025).

Rooting strategies and interspecies competition strongly modulate tree water uptake dynamics (Berkelhammer *et al*., 2020; Hackmann *et al*., 2025). In mixed stands, spruce typically maintains a shallow rooting system, relying primarily on near-surface soil moisture (Brinkmann *et al*., 2019) and becoming highly vulnerable to summer droughts when the topsoil dries out (Bishop & Dambrine, 1995; Schuldt *et al*., 2020; Grams *et al*., 2021). In contrast, deciduous species such as *Fagus sylvatica* (referred to as “beech” hereafter) can better adapt to reductions in topsoil moisture due to their deeper rooting systems in mixed stands, allowing access to deeper soil water (Brinkmann *et al*., 2019; Meusburger *et al*., 2022; Hackmann *et al*., 2025). Moreover, soil texture appears to play only a limited role in determining depth of tree water uptake (Hackmann *et al*., 2025). It remains unclear, however, whether spruce in mono-specific natural forests can develop deeper roots and exhibit more flexible water-use strategies, as most isotope-based studies have been conducted in mixed stands, where competitive interactions dominate (Goldsmith *et al*., 2019; Zhang *et al*., 2021; Grams *et al*., 2021; Gessler *et al*., 2022; Muhic *et al*., 2023; Floriancic *et al*., 2024; Hackmann *et al*., 2025).

Recent isotope-based ecohydrological studies have emphasized that tree water uptake depth is strongly influenced by atmospheric wetness and soil water turnover, quantifying how quickly precipitation replenishes the soil. In an extremely wet montane forest in the Southeast Andes, with annual precipitation exceeding 4000 mm, Burt et al., (2023) showed shallow root water uptake across multiple tree species, with xylem isotopes resembling wet-season precipitation during the wet season and dry-season precipitation during the dry season. Similarly, Zuecco et al., (2026) reported high soil water turnover in a wet headwater catchment in the Italian pre-Alps, where beech and chestnut trees primarily used summer precipitation during the growing season. In the boreal forests of Finnish Lapland, Muhic et al., (2023) showed rapid transmission of rainfall and snowmelt through the soil profile. Subsequent isotopic labelling experiments at their site revealed fast vertical propagation of labelled water through the soil, with spruce trees taking up this water within 30 hours after the start of irrigation (Muhic *et al*., 2024). Multiple spatial snapshot studies across Switzerland have likewise shown that trees in wetter regions predominantly use soil water recharged by recent precipitation (Allen *et al*., 2019; Goldsmith *et al*., 2022; Floriancic *et al*., 2025; Burt *et al*., 2025). These findings suggest that tree water uptake dynamics are – besides biological factors – strongly influenced by the timing of soil water replenishment, which remains understudied in high-elevation spruce forests.

There is a clear need to characterize how spruce in subalpine mono-specific forests responds to seasonal water availability and soil recharge dynamics. In this study, we investigated seasonal shifts in water sources used by spruce trees in the Davos Seehornwald forest in eastern Switzerland across three consecutive growing seasons (2020–2022), including the 2022 drought year. Using stable water isotopes (δ¹⁸O, δ²H) in precipitation, bulk soil water, and xylem water, we quantified how much spruce water uptake was from deeper (50-70 cm) and shallower soil layers (0-10 cm), and how spruce water uptake transitions from cold– to warm-season-derived precipitation, and related these patterns to seasonal soil water recharge. We further assessed whether spruce in a mono-specific natural forest accessed soil water deeper than the first 30 cm, and examined how the timing of seasonal water sources related to surface energy availability and soil water recharge.

## Materials and Methods

The study was conducted in a subalpine evergreen needleleaf forest located in the eastern Swiss Alps (Davos Seehornwald; CH-Dav; 46°48’55.2” N, 9°51’21.3”, 1639 m a.s.l.; Figure 1). CH-Dav is an ICOS (Integrated Carbon Observation System) Class 1 Ecosystem station, dominated by Norway spruce (*Picea abies* (L.) Karst), with average tree height of 17.5 m, maximum height of 41 m, and average tree age of 120 years, with the oldest tree estimated to be 350 years old. The forest has a patchy understory composed of blueberry (*Vaccinium myrtillus* and *Vaccinium gaulterioides*) and mosses (*Sphagnum* sp. Ehrh. and *Hylocomium splendens*), covering about 30% of the forest floor (Krebs *et al*., 2024). The region experiences a temperate climate, with mean annual precipitation of ∼876 mm and mean annual temperature of ∼4.3 °C (1997-2024). The site features Chromic Cambisols and Rustic Podzols soils, with soil texture ranging from sand to sandy loam (Jörg, 2008; Krebs *et al*., 2025).

**Figure 1:**
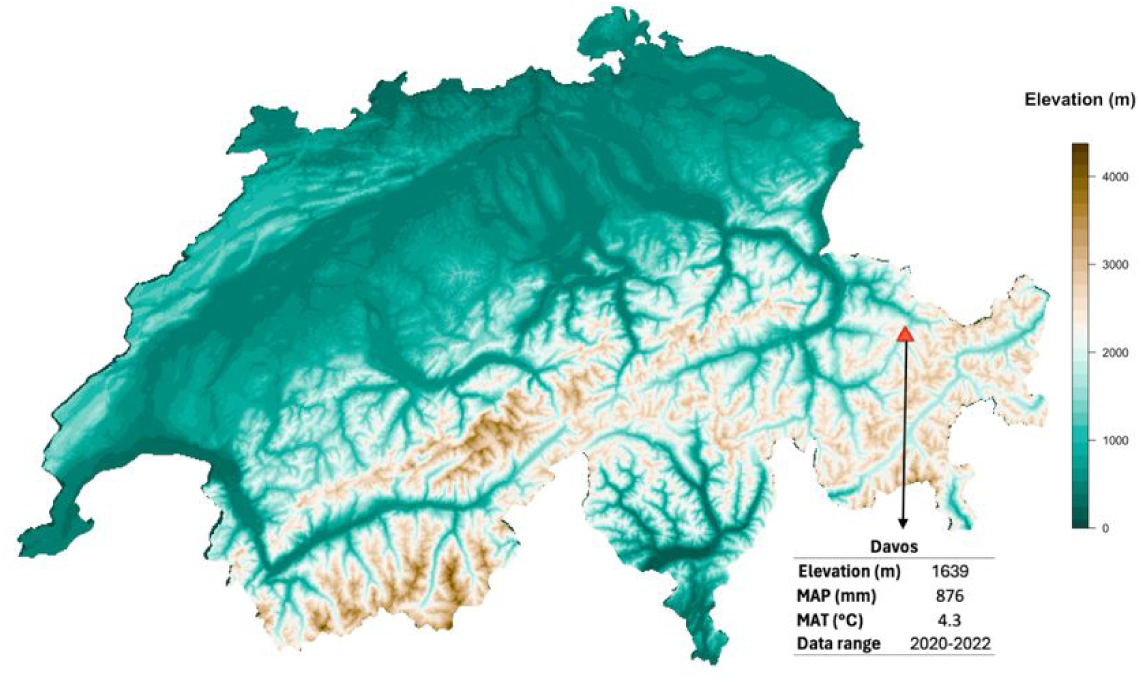
Location of the Davos Seehornwald spruce forest (CH-Dav) in the eastern Swiss Alps. The map shows the site overlaid on a digital elevation model (SwissTopo) and indicates key meteorological conditions. CH-Dav is a subalpine evergreen needleleaf forest and a long-term ecological monitoring site.

At CH-Dav, water vapor fluxes have been continuously measured since 1997 using the eddy covariance (EC) technique to estimate ecosystem-scale evapotranspiration fluxes (Feigenwinter *et al*., 2025). The EC system is installed at 35 m above ground and consists of a three-dimensional sonic anemometer for wind speed measurements and enclosed-path infrared gas analyzers (IRGA) for measuring dry mole fractions of water vapor. Half-hourly H_2_O fluxes (FW, mmol H_2_O m^−2^ s^−1^) were calculated by maximizing the covariance between turbulent vertical wind speed and water vapor concentration, applying a lag search of 0-10 s combined with predefined nominal time lags using the EddyPro v7 software (LI-COR Inc., USA). Details on flux processing and evapotranspiration calculations are provided in Shekhar et al., (2024). Potential evapotranspiration (PET) was calculated using the Thornthwaite method (Thornthwaite, 1948; Oudin *et al*., 2005). The site is also equipped with soil water content sensors (Decagon ECH_2_O EC-20, Pullman, WA, USA). For this study, we used soil moisture measurement at 15 cm depth from 2020 to 2022, which provided the most continuous record with minimal data gaps.

## Sampling and isotopic analyses

Bulk soil and xylem samples from 10-16 spruce trees were collected approximately biweekly from May to September during 2020–2022. Soil was sampled in 10-cm increments from the surface to 70 cm using a Pürkhauer soil corer with a diameter of 28 mm (Goecke, Schwelm, Germany). In 2020, the uppermost soil layers were sampled separately at 0-5 and 5-10 cm, whereas in 2021 and 2022, sampling started at 0-10 cm. Subsequent sampling was taken in 10-cm intervals down to 60-70cm. Soil samples were stored in 12 ml airtight glass exetainers (Labco Ltd., UK) and stored on ice in the field until placed in a freezer at –23 °C. Xylem samples were obtained from branches of mature spruce trees from 10-12 m height. The bark and phloem were manually removed from twig samples, and the remaining xylem was quickly placed in 12 ml airtight glass exetainers, similar as the soil samples.

We extracted water from soil and xylem samples using a cryogenic water extraction line (Ehleringer *et al*., 2000). Samples in the 12 ml glass exetainers were immersed in an 80 °C water bath. The evaporated water was collected in U-tubes submerged in liquid nitrogen (–196 °C) for condensation. Extraction was conducted under a vacuum of approximately 0.03 hPa for 2 hours. After extraction, the U-tubes were sealed and allowed to thaw. The condensed water was collected and stored in 1.5 ml airtight glass vials (Macherey-Nagel, Düren, Germany) until stable water isotope analysis.

Isotope analysis of δ^18^O and δ^2^H was performed using a high-temperature conversion/elemental analyzer (TC/EA; Finnigan MAT, Bremen, Germany) coupled to a DeltaplusXP isotope ratio mass spectrometer via a ConFlo III interface (Finnigan MAT; Werner et al., (1999)), employing the carbon reduction method of Gehre et al., (2004). Water samples were automatically injected into a gas chromatography PAL autosampler (CTC, Zwingen, Switzerland). To ensure consistency, we adhered to the approach of Werner & Brand (2001) for sample and laboratory standard positioning and identical treatment during measurements. Post-run corrections (offset, memory effects, drift) were also performed as described in Werner & Brand (2001). Long-term quality control using lab standard water (WP-0503-Z0010B) yielded precisions of ±0.17‰ for δ^18^O and ±0.48‰ for δ^2^H. All samples were calibrated against the Vienna Standard Mean Ocean Water (V-SMOW) and Standard Light Antarctic Precipitation (SLAP).

As cryogenic vacuum extraction (CVD) can introduce systematic biases in δ²H measurements (Chen *et al*., 2021; Allen & Kirchner, 2022), δ¹⁸O was used as the primary tracer for source attribution and mixing analysis. While the suitability of CVD for soil and xylem water extraction continues to be debated, it remains the most widely applied method in ecohydrological studies (Ceperley *et al*., 2024). Additionally, several ecohydrological studies have shown that trees predominantly use tightly bound soil water rather than mobile soil water (Brooks *et al*., 2010; Berghuijs & Allen, 2019; Sprenger & Allen, 2020), supporting the use of bulk soil water samples as representative of the water pool accessible to roots.

## Bayesian mixing model (HydroMix)

To estimate the seasonal origin of tree water, we use HydroMix, a Bayesian mixing model specifically designed for ecohydrological applications with limited tracer data (Beria *et al*., 2020). Traditional Bayesian mixing models infer mixing ratios by fitting probability density functions (pdfs) to stable water isotope ratios in different sources (e.g., precipitation, soil water) and mixtures (e.g., groundwater, xylem water), and then estimating the posterior distribution of mixing ratios using standard Bayesian inference principles (Parnell *et al*., 2010; Popp *et al*., 2025). However, sparse sampling (<20-30 measurements per day and location) often makes it difficult to robustly estimate source and mixture pdfs with limited isotopic data. HydroMix overcomes this limitation by adopting a bootstrap approach using all possible combinations of observed source and mixture isotopic ratios, and a given distribution of mixing ratios to simulate mixture isotopic ratio in a Monte Carlo setup. HydroMix then optimizes a likelihood function based on the mismatch between the simulated and observed mixture isotopic ratio, thereby bypassing the need for an analytical pdf for the source and mixture isotopic ratios. This approach has been shown to perform well in synthetic benchmarking experiments (Beria *et al*., 2020) and in real-world applications (Beria *et al*., 2020; Sprenger *et al*., 2024).

Here, we apply HydroMix to estimate (1) the relative contribution of shallow (0-10 cm) and deep (50-70 cm) soil water to xylem water, and (2) the relative contribution of warm-season (May-September) and cold-season (October-April) precipitation to tree water uptake across successive growing seasons (2020-2022). HydroMix was run using δ¹⁸O as the primary tracer, and the mean and uncertainty (±1 standard deviation) of mixing fractions were derived from the top 20% highest-likelihood simulations.

## Results

### Soil water and xylem isotope ratios over time

The dual isotope plots (Figures 2, S1, S2) showed clear depth-dependent patterns in soil water, with shallow soil layers being consistently more enriched in heavier isotopes than deeper soil layers during most of the growing season. The soil water lines, derived from soil samples from all depths and all locations on each sampling day, had slopes ranging from 4.67 to 7.81. These slopes were generally lower than the slope of the Local Meteoric Water Line (LMWL; slope=7.48; Figure S3), indicating the effect of kinetic fractionation resulting from evaporation from the upper 0-10 cm soil layer (Beria *et al*., 2018). The LMWL slope itself (7.48) was slightly lower than the Global Meteoric Water Line (GMWL; slope=8), but consistent with previous studies from the European Alps (Michelon *et al*., 2023; Zuecco *et al*., 2026). In deeper soil layers (50-70 cm), evaporative signatures were largely absent, except on 18 September 2020 (Figure 2), when the observed enrichment likely reflected downward infiltration of evaporatively enriched shallow soil water.

**Figure 2:**
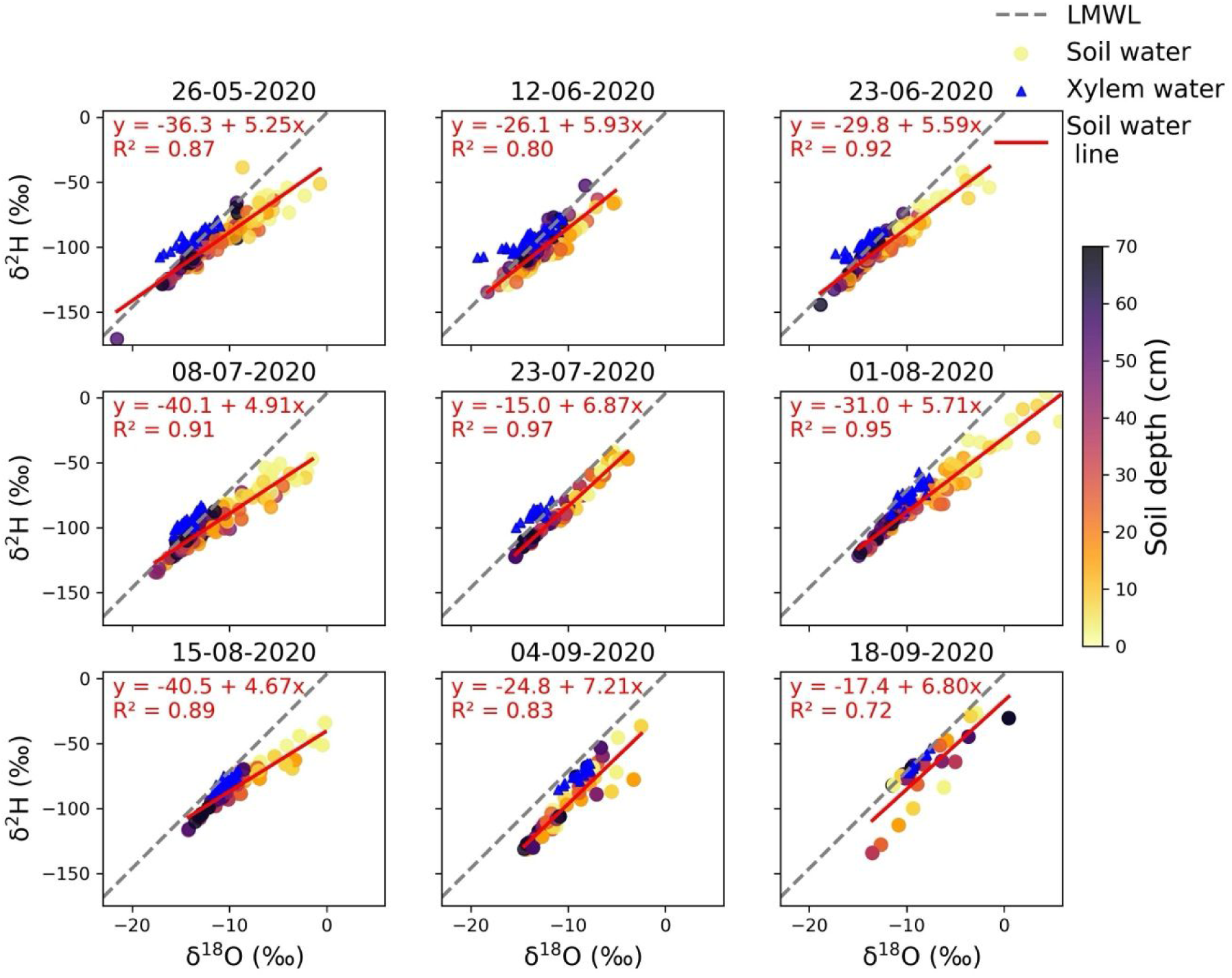
Dual isotope relationships (δ¹⁸O and δ²H) for bulk soil water and xylem water for the 2020 growing season in the Davos Seehornwald forest (CH-Dav), dominated by spruce. Soil water isotopes are plotted as colored dots (colors indicating soil depth) and xylem water isotopes are shown in blue triangles. Local Meteoric Water Line (LMWL) is derived from precipitation samples collected at the site during 2020 to 2022 (see Figure S3) and is shown as dashed grey line in each subplot. Dual isotope relationships for the 2021 and 2022 growing seasons are shown in Supporting Information (Figures S1, S2).

Soil δ¹⁸O values showed a strong vertical gradient from May to August in all years (Figures 3, S4, S5), with δ¹⁸O increasing non-linearly toward the shallowest soil layers. As the growing season advanced, deeper soil layers became increasingly enriched in heavier isotopes, reflecting the downward infiltration of isotopically enriched growing-season rainfall and resulting in vertical homogenization of soil water isotopes.

**Figure 3:**
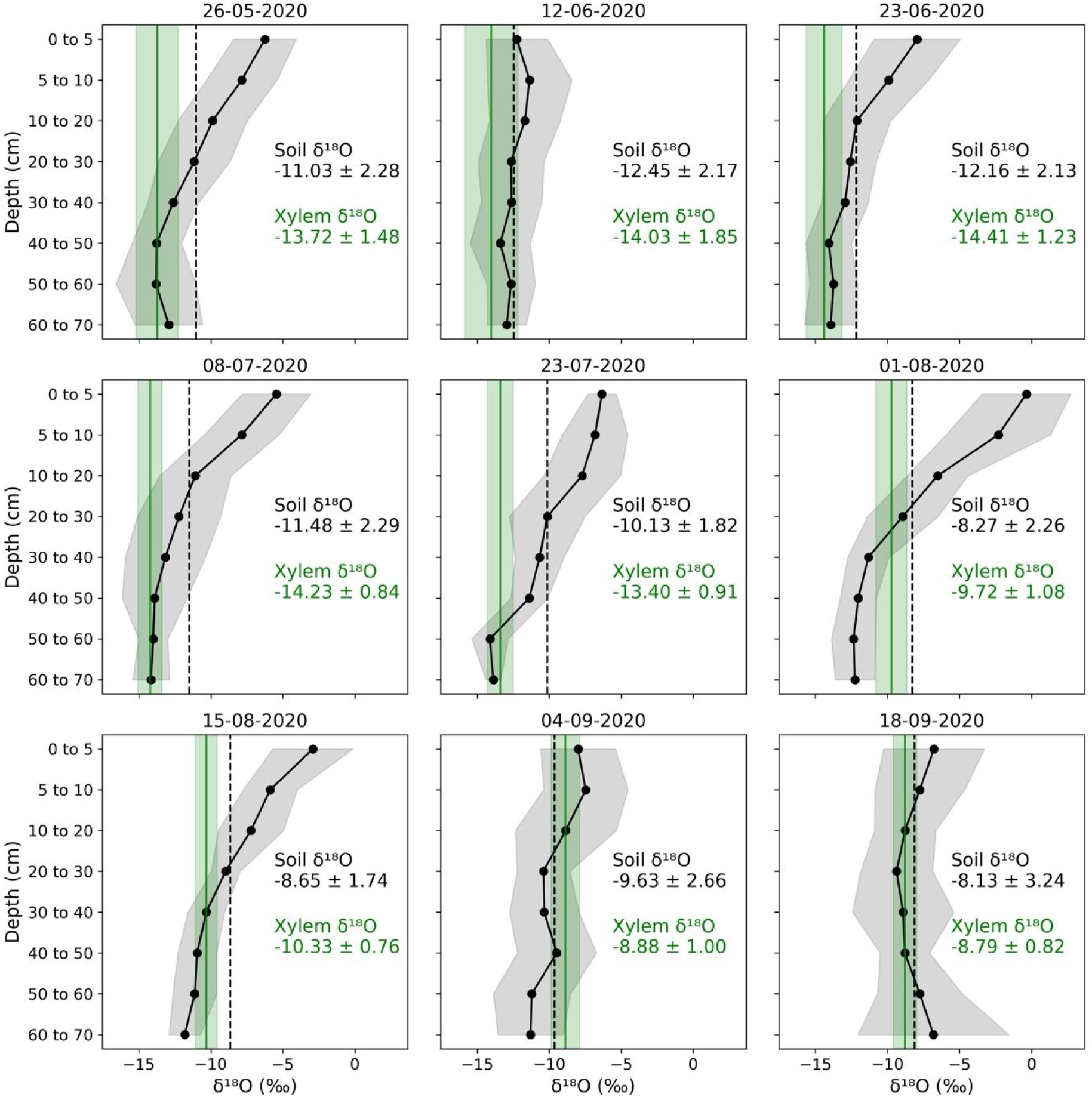
Vertical soil water and xylem water isotopic profiles (δ¹⁸O) averaged across sampling locations in 2020 (corresponding figures for 2021 and 2022 are shown in Supplementary Figures S4 and S5). Each panel shows the mean ± standard deviation of δ¹⁸O values in xylem water (green band) and in soil water (gray band) at multiple depths for each sampling date, calculated from the individual sampling points. The green line represents the daily mean xylem δ¹⁸O values across the sampled trees, and the black dotted line depicts the soil δ¹⁸O values averaged across all depths and sampled locations for each day.

Across all years, xylem δ¹⁸O values increased over the course of the growing season (Figure 4). Early in the season, xylem waters were relatively depleted in heavier isotopes, followed by gradual enrichment toward late summer. The timing of enrichment in xylem water differed among years with 2022, the driest of the three years, showing the earliest shift toward enriched δ¹⁸O values. Overall, both deep soil (50-70 cm) and xylem water displayed a consistent seasonal progression toward more enriched isotopic compositions from spring to late summer. A detailed qualitative description of intra-annual variability in soil and xylem isotopic dynamics is provided in the Supplementary Information (see Notes S1).

**Figure 4:**
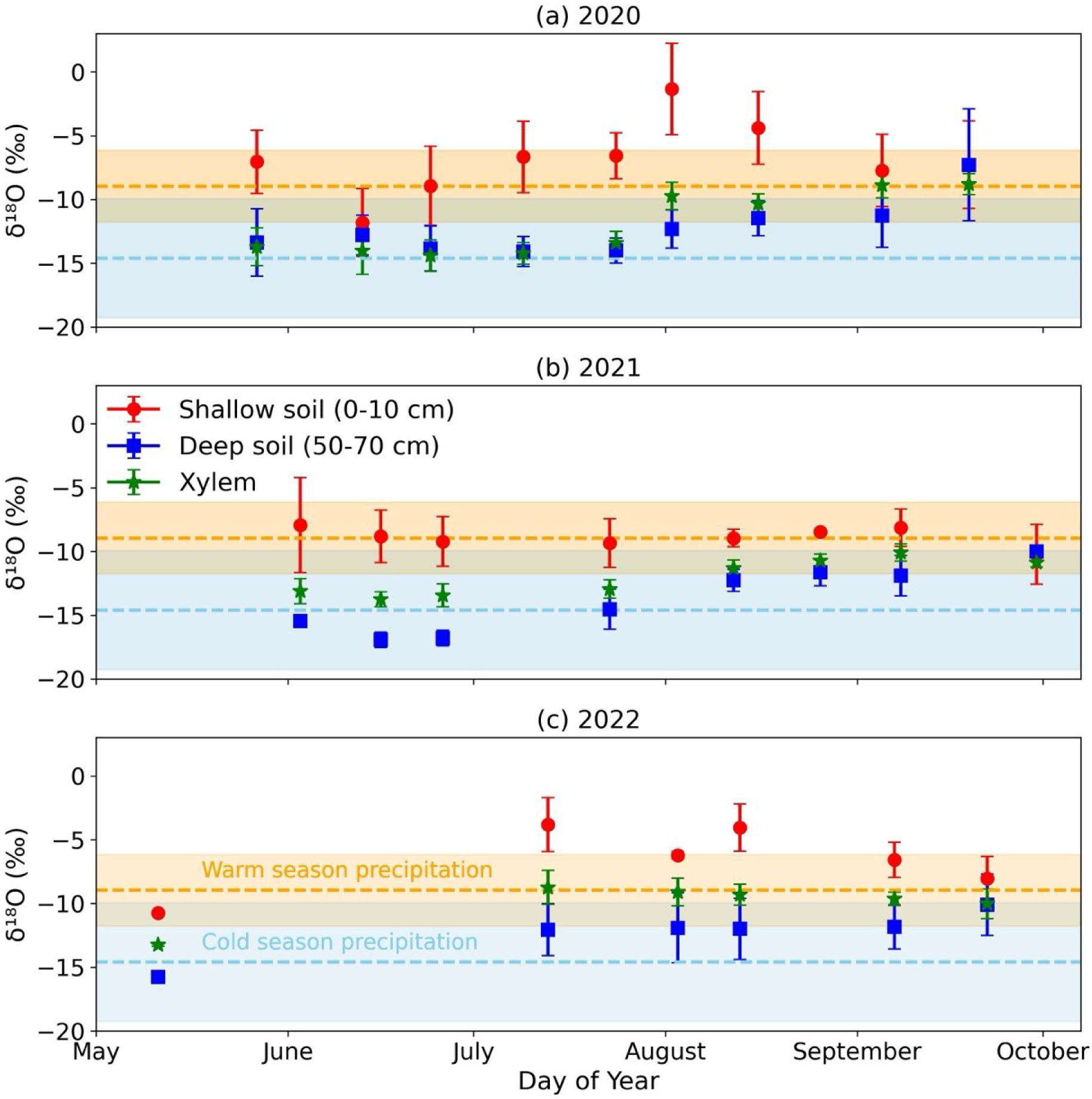
Temporal development of shallow (0-10 cm), deep soil (50-70 cm), and xylem δ¹⁸O in (a) 2020, (b) 2021, and (c) 2022. Each panel shows the mean ± standard deviation of isotope ratios in shallow and deep soil water and in xylem water, averaged across all sampling locations for each sampling date. The orange and blue bands show the mean ± standard deviation of warm-season (May-September) and cold-season (October-April) precipitation δ¹⁸O values, with dashed lines showing average δ¹⁸O values, calculated from precipitation samples collected between 2020-2022.

### Deep soil water (50-70 cm) as the dominant contributor to root water uptake

Readers are reminded that we only considered two sources (0-10 cm for shallow soil water and 50-70 cm for deep soil water) as their isotopic ratios were clearly distinct (Figure 4). Across the three years, deep soil water (50-70 cm) contributed more than 50% to spruce water uptake for most periods (Figure 5a). In 2020, more than 80% tree water uptake came from deeper soil water during spring and early-summer, transitioning towards shallower soil water as the growing season advanced. In 2021 and 2022, both shallow and deep soil water contributions were comparable throughout the growing season, with higher contributions from deeper soil water in 2021, and higher contributions from shallower soil water in 2022. This contrasts with findings from mixed forests, where spruce typically relies on near-surface soil layers and shows limited access to deeper soil water (Bolte & Villanueva, 2006; Grams *et al*., 2021; Hackmann *et al*., 2025).

**Figure 5:**
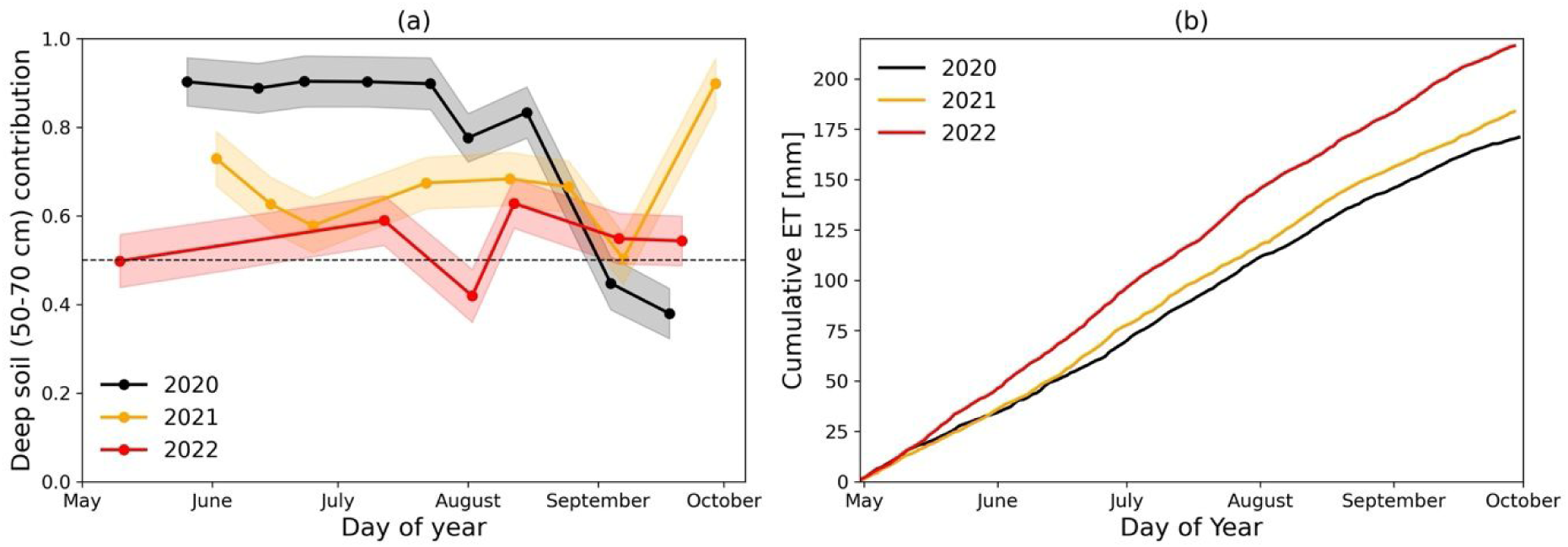
(a) Deep soil water (50-70 cm) contributions to xylem water, estimated using δ¹⁸O for different sampling dates in 2020 (black), 2021 (yellow), and 2022 (red). Colored dots show the mean of the posterior distribution derived from the top 20% highest-likelihood simulations, and the shaded envelope represents ±1 standard deviation. (b) Cumulative actual evapotranspiration from 1 May to 30 September for 2020 – 2022.

A pronounced deep-to-shallow seasonal shift was evident in late 2020 (Figure 5a). Early in the growing season, mean xylem δ¹⁸O values matched with 50-70 cm soil δ¹⁸O values (Figure 3), indicating dominant deep uptake during May and June when atmospheric demand and actual evapotranspiration were low (Figure 5b). As the season progressed and evaporative demand increased, spruce shifted its water uptake toward shallower soil waters, with shallow soil (0-10 cm) contributions increasing through August. In contrast, 2021 and 2022 exhibit more balanced contributions from shallow and deep soil water throughout the season (Figure 5a), reflecting year-to-year variability in water uptake strategies. In 2022, exceptionally high evaporative demand and rapid increases in cumulative evapotranspiration (Figure 5b) coincided with higher uptake from shallower soil sources, showing spruce’s reliance on shallower water pools to sustain periods of higher water demand (Figure 5a).

### Contributions of cold-season vs. warm-season precipitation to tree water uptake

Parallel to the transitions from deep to shallow water uptake, Bayesian mixing analysis during the three growing seasons revealed a consistent transition from cold– to warm-season precipitation as the dominant source of spruce water uptake (Figure 6). Early in each growing season, xylem water uptake showed preferential uptake from deeper soil water pools recharged primarily from cold-season precipitation (Figures 5a, 6), followed by a gradual shift toward warm-season precipitation.

**Figure 6:**
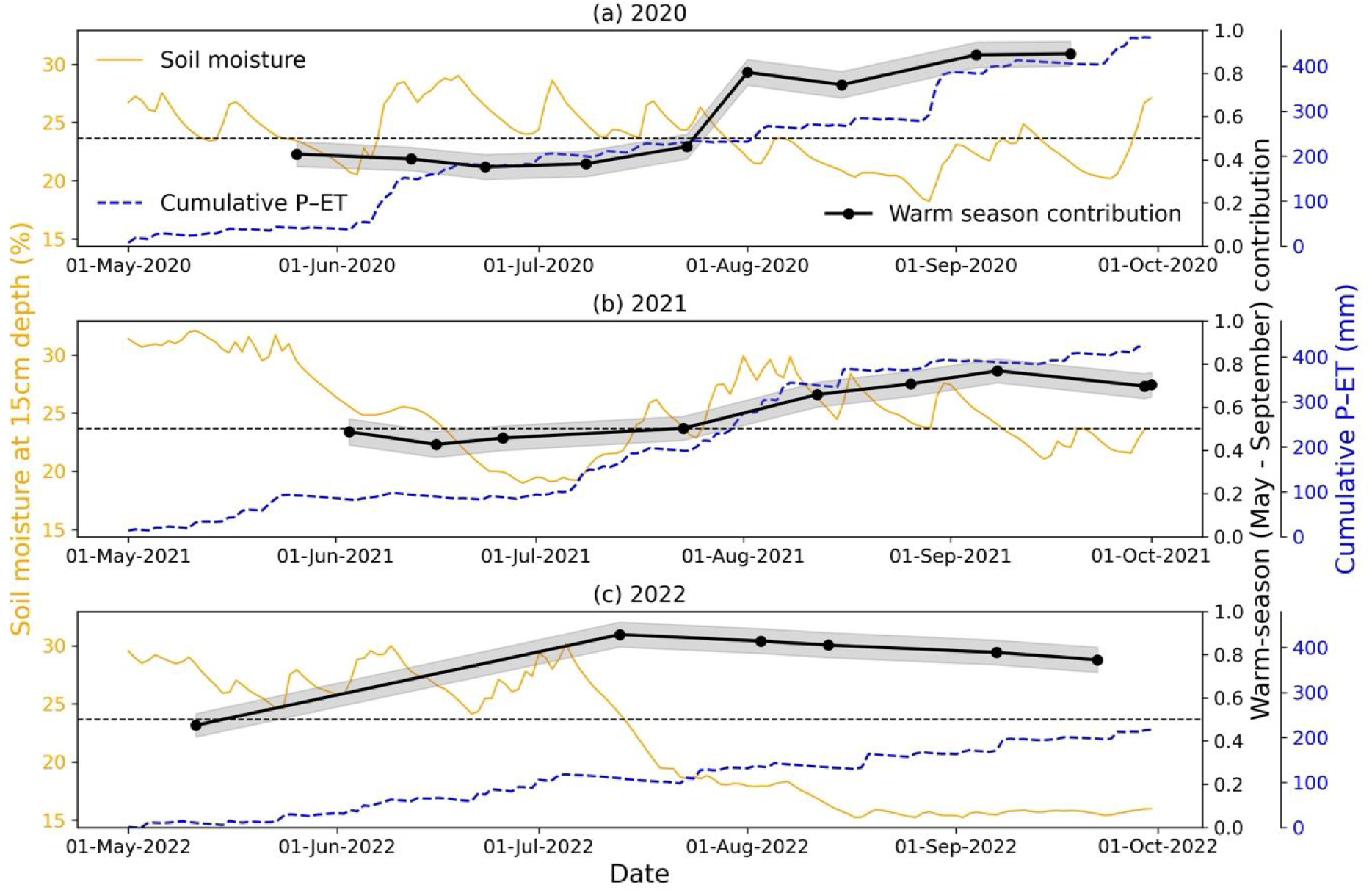
Warm season (May-September) precipitation contribution to xylem water (estimated using δ¹⁸O, black line) shown in relation to soil moisture at 15cm depth (yellow line) and cumulative growing-season soil water recharge (precipitation minus actual evapotranspiration; blue dashed line) for (a) 2020, (b) 2021, and (c) 2022.

In 2020, cold-season contribution (defined as 1 – warm-season contribution) remained around ∼60% from May through July, primarily originating from deeper soil layers (>40 cm; also see Notes S1 in Supplementary Information), followed by a distinct shift toward warm-season precipitation in August, contributing 80-90% to tree water uptake (Figure 6). In 2021, June and July spruce water uptake showed a slightly greater contribution from warm-season precipitation (∼50%) than in June and July 2020, and by August 2021, warm-season contribution approached ∼80%, similar to the 2020 dynamics. In 2022, only one early growing-season sampling campaign was conducted in May, when xylem water reflected an approximately equal mixture of cold– and warm-season precipitation (Figure 6). By mid-July 2022, spruce had already shifted to more shallower soil layers predominantly recharged from warm-season precipitation (>85% contribution) and maintained this reliance through late summer.

The earlier transition from cold– to warm-season precipitation in 2022 is consistent with its exceptionally warm and dry conditions. These conditions increased seasonal energy availability, resulting in higher cumulative growing-season evapotranspiration (216 mm in 2022; Figure 5b) relative to 2020 (171 mm) and 2021 (184 mm). Additionally, earlier peak energy availability also likely advanced phenological development. Interestingly, during 2022, cumulative evapotranspiration also increased much faster than in the other years (Figure 5b), suggesting that atmospheric energy availability may have been the primary driver for tree water uptake.

### Soil water recharge as the dominant control on seasonal water uptake

Across the three years, the transition of spruce water uptake from cold-season to warm-season precipitation did not coincide with any specific 15-cm soil moisture threshold (Figure 6). In 2020, the shift occurred in late July when 15-cm soil moisture dropped below ∼24% (Figure 6a), but in 2021, a similar timing was seen despite much wetter upper soils in August, when over 60% of water uptake reflected warm-season precipitation (Figure 6b). In 2022, a decline in 15-cm soil moisture was loosely aligned with the water uptake transition to warm-season precipitation, but the limited early-season sampling in 2022 prevents a definitive interpretation (Figure 6c). Together, these observations indicate that soil moisture in shallow soil does not regulate when spruce switches from cold– to warm-season precipitation.

In contrast, the timing of the seasonal transition aligned with growing-season soil water recharge (Figure 6). Across 2020-2022, cumulative precipitation minus actual evapotranspiration remained positive throughout the growing season, indicating that precipitation inputs consistently exceeded evaporative losses and recharged the soil column – even in 2022, the driest of the three years, when net soil recharge from May-September 2022 exceeded 200 mm. This newly infiltrated water was consistently taken up by spruce, indicating that recharge-driven replenishment of the soil profile plays a central role in enabling seasonal shifts toward warm-season precipitation.

Evapotranspiration patterns further revealed that atmospheric energy availability (through enhanced shortwave net radiation), rather than soil moisture, constrained ecosystem-scale water use in Davos, Seehornwald. Across the three seasons, May to September precipitation and actual evapotranspiration were inversely related, whereas actual evapotranspiration and potential evapotranspiration showed a positive relationship. For example, 2022 had the lowest growing-season (May-September) precipitation (434 mm) but the highest actual evapotranspiration (216 mm), coinciding with the highest potential evapotranspiration (1044 mm). Conversely, 2020, the wettest growing-season (635 mm), had the lowest actual evapotranspiration (171 mm) and the lowest potential evapotranspiration (927 mm). 2021 was a moderately wet growing-season with 606 mm precipitation, 184 mm actual evapotranspiration, and 941 mm potential evapotranspiration. This suggests that atmospheric energy availability, as depicted by potential evapotranspiration, was the main driver of actual evapotranspiration, with soil wetness playing a limited role. This behavior, seen particularly in energy-limited ecosystems, has been documented as “drought paradox”, where higher actual evapotranspiration occurs during warm, dry years despite reduced soil moisture, and has been shown in many regions including Europe (Teuling *et al*., 2013; Orth & Destouni, 2018; Massari *et al*., 2022), southeastern USA (McQuillan *et al*., 2022), the Canadian Rockies (Langs *et al*., 2021), the Pyrenees (Vicente-Serrano *et al*., 2021), and southeastern Australia (Stephens *et al*., 2023).

## Discussion

### Deep soil water uptake by Norway spruce in a mono-specific natural forest

In mixed stands, spruce fine roots are typically concentrated in the upper soil layers (Bolte & Villanueva, 2006; Grams *et al*., 2021). Isotope-based studies conducted in mixed forests have inferred that spruce water uptake happens primarily from the 10-30 cm soil depth range (Brinkmann et al., 2019; Floriancic et al., 2024; Hackmann et al., 2025; Zhang et al., 2021). These results have contributed to the widespread perception that spruce is highly vulnerable to drought because of topsoil drying (Schuldt *et al*., 2020; Grams *et al*., 2021). However, this view largely stems from studies in mixed stands, where spruce typically competes with deeper-rooted species such as beech, potentially constraining its effective rooting depth.

Our isotope-based analyses in the mono-specific natural forest at Davos revealed substantial and persistent spruce water uptake from deeper soil layers (50-70 cm; Figure 5a), challenging the prevailing view of Norway spruce as a predominantly shallow-rooted and drought-sensitive species. In early 2020, δ¹⁸O reflected contributions from deep soil water, with up to ∼90% of tree water uptake derived from deeper soil water pools. This underscores the importance of winter and spring precipitation, and subsequent snowmelt, in recharging deeper soil reservoirs that sustain early-season vegetation growth. In the European Alps, this period is undergoing rapid changes due to climate warming, including increasing winter and spring precipitation, instead of winter snowfall, and earlier snowmelt, with important implications for soil water recharge (Scherrer *et al*., 2025).

Across the three years, we observed interannual variability in the relative contributions from shallow and deep soil water. In 2020, spruce showed a clear transition from deep to shallower soil water uptake as the growing season progressed. This pattern is consistent with observations from conifer species in the Rocky Mountains (Berkelhammer *et al*., 2020), where high-growth periods were associated with reliance on shallow, recently infiltrated water. In 2021, deep soil water contribution exceeded 50% for most of the growing season, whereas in the dry and warm year of 2022, shallow soil contribution was slightly higher than in the other years, showing plasticity in water uptake strategies by spruce trees. These differences reflect year-to-year variability in atmospheric demand and timing of soil water recharge and point to dynamic adjustment of uptake strategies under contrasting hydroclimatic conditions.

Overall, our results indicate that in the absence of interspecific competition, spruce rooting space is not tightly constrained, allowing water uptake along the entire soil profile, including deep soil layer. These results highlight that spruce water-use strategies in humid, mono-specific natural forests are more flexible than suggested by studies from drier or mixed-species forests. Deep soil water emerges as a vital and resilient reservoir for spruce in subalpine forests, even during warm and dry years. The plasticity in spruce water uptake provides rare empirical evidence of greater rooting depths and adaptive water-use strategies in spruce forests. The Davos Seehornwald observations thus offer an important foundation for future root water uptake modeling studies and for refining conceptual models of spruce ecohydrology under changing hydroclimatic regimes.

### Seasonal patterns in soil and xylem isotopes and the underlying mechanisms

Our observations revealed consistent seasonal dynamics in soil and xylem isotopes that extend existing evidence of evaporative enrichment in topsoil (Muhic *et al*., 2023) and vertical mixing in forest soils (Brinkmann *et al*., 2019). Across all three years (2020-2022), shallow soils (0-10 cm) remained isotopically enriched in heavier isotopes relative to deeper layers, while δ¹⁸O values in deeper soil progressively increased over the growing season (Figure 7). This pattern indicates the downward transmission of isotopically enriched summer rainfall and increasing vertical homogenization of the soil profile. Similar seasonal enrichment of deeper soil layers has been reported in boreal and Swiss forests (Brinkmann *et al*., 2019; Muhic *et al*., 2023), where rapid soil water turnover facilitates vertical mixing.

**Figure 7:**
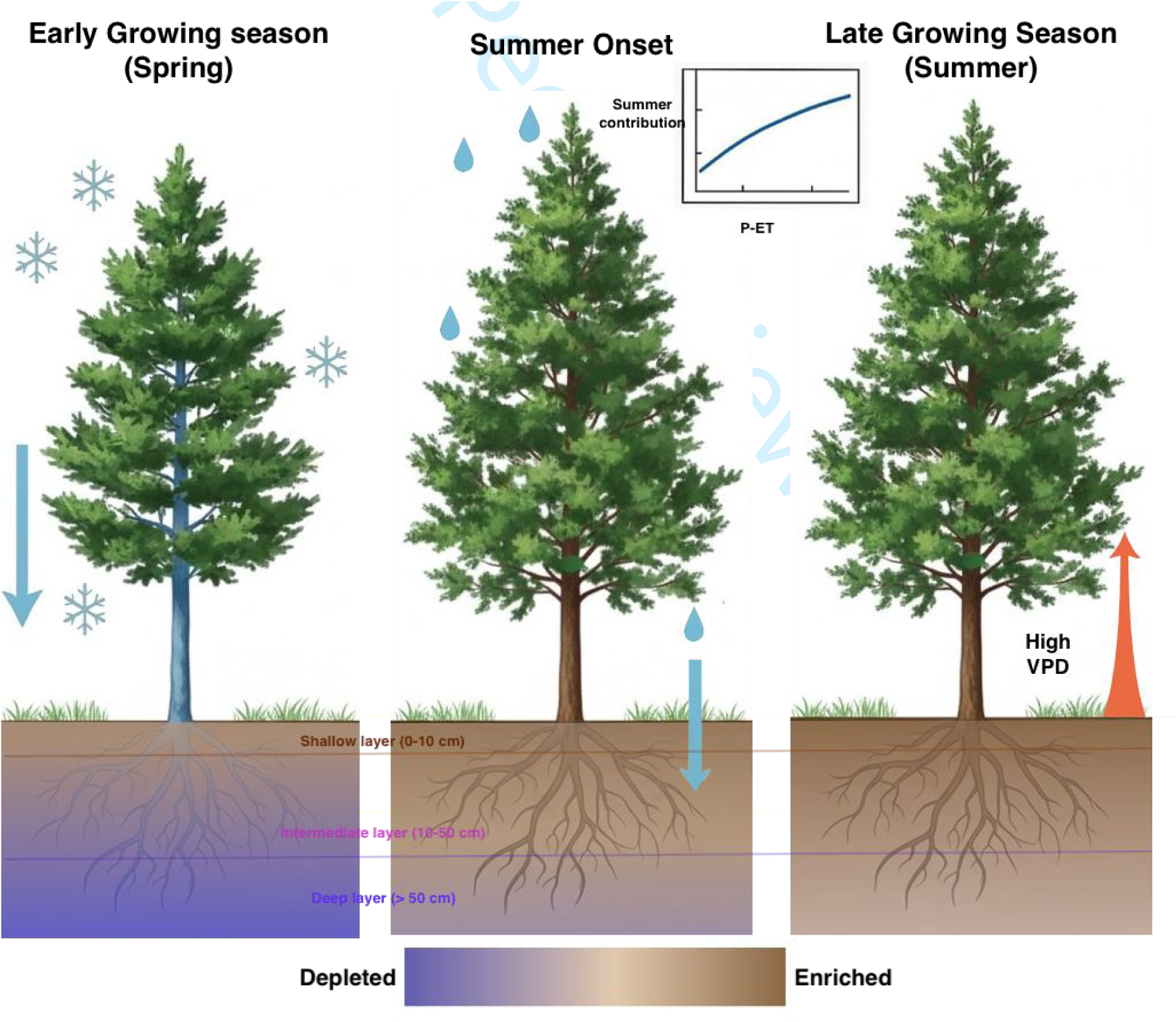
Conceptual illustration of seasonal soil plant dynamics in the Davos Seehornwald spruce stand. Early in the growing season (left), spruce predominantly accesses deeper soil water (>50 cm) recharged by cold-season precipitation, while shallow soils remain isotopically enriched. As the growing season progresses (middle), summer precipitation transmits isotopically enriched waters from shallow soil to deeper soil layers. Under warm and dry conditions with high atmospheric demand (right), spruce increasingly relies on recently recharged growing-season precipitation, drawn from both shallow and deep soils, with water uptake strongly coupled to soil recharge and vapor pressure deficit (VPD). Shading in the soil layers and in the tree bark indicates relative isotopic ratios, with blue shades representing cold-season depleted isotopic ratios and brown shades depicting warm-season enriched isotopic ratios. The arrows denote dominant water fluxes and atmospheric controls.

Xylem δ¹⁸O values mirrored deeper soil dynamics, becoming progressively enriched over the growing season and reflecting a clear shift in spruce water uptake toward summer precipitation signature in shallow soil depths. These seasonal uptake dynamics were consistent with observations from other humid forest ecosystems, where conifers increasingly utilize summer precipitation as the growing-season progresses (Berkelhammer *et al*., 2020; Zuecco *et al*., 2026). The closest comparable study came from the Rocky Mountains of North America, where conifer trees showed increased reliance on summer precipitation during high-growth years, hypothesized as seasonal shallowing of the effective root uptake depth (Berkelhammer *et al*., 2020), comparable to the early and pronounced shift in seasonal uptake with higher contributions from shallower soil layers at Davos Seehornwald in 2022 (Figures 5a, 6c). In boreal Finland, spruce incorporated newly infiltrated water within 30 hours following irrigation (Muhic *et al*., 2024), demonstrating their capacity for rapid uptake of recent precipitation from the top soil. Similarly, in Central Europe, water consumption by another evergreen coniferous species (Douglas fir) was closely linked to dynamics of the organic surface layer (Hackmann *et al*., 2025), with trees rapidly capturing small rainfall inputs during periods of high demand (Dietrich & Kahmen, 2019).

The transition in spruce water uptake from cold-season to warm-season precipitation at Davos Seehornwald appeared to be primarily controlled by positive soil recharge (cumulative precipitation minus evapotranspiration since May 1) and atmospheric demand, rather than by 15-cm soil moisture or fixed rooting constraints. The role of positive soil recharge is evident across all years (Figure 6) and is particularly exemplified in 2021, when substantial late-July soil recharge coincided with a sustained shift of water uptake toward warm-season precipitation (Figure 6b). Atmospheric control becomes more apparent in the exceptionally warm and dry year of 2022, when the shift from cold– to warm-season precipitation occurred markedly earlier than in 2020 and 2021 (Figure 6c).

Together with recent findings from other wet and snow-influenced catchments (Burt *et al*., 2023; Brighenti *et al*., 2024; Zuecco *et al*., 2026), our study supports an emerging view of dynamic coupling between soil water recharge, atmospheric energy availability, and tree water use in humid forests. In such landscapes, growing-season precipitation continuously replenishes soil water and infiltrates to deeper depths. Under high radiative and evaporative demand, this newly infiltrated water is rapidly taken up by trees from different soil depths, including >50% uptake from deeper layers to sustain ecosystem evapotranspiration.

## Conclusions and outlook

Isotopic and hydrometeorological evidence from three continuous growing seasons in the Davos Seehornwald mono-specific natural forest revealed pronounced plasticity in spruce water uptake, including consistent access to deep soil water (50-70 cm). These findings challenge the prevailing view of Norway spruce as a shallow-rooted and drought-specific species, instead indicating that spruce water use strategies are strongly conditioned by interspecies competition. Consequently, drought sensitivity inferred primarily from mixed, low-elevation forests does not necessarily apply to wide-spread spruce natural forests at higher elevations.

At higher elevations, where conifer forests prevail and spruce often coexists with deep-rooted species such as larch, competitive constraints on spruce may be relaxed, allowing greater plasticity in water uptake. As climate change continues to lengthen growing seasons and increase energy availability in mountain regions, this flexibility may enable spruce forests to sustain or even increase evapotranspiration, as seen during the 2022 drought year. However, the increasing reliance of mono-specific natural spruce stands to summer-rainfall-recharged soil water may also increase their vulnerability to more frequent droughts and heat waves, specifically in scenarios when cumulative evapotranspiration exceeds cumulative precipitation over a sustained period. Extending similar ecohydrological investigations to other regions will be critical for untangling these mechanisms and for assessing the broader relevance of adaptive strategies and their implications for forest resilience under future climates.

More broadly, our results demonstrate that tree water uptake strategies cannot be inferred from rooting depth distributions or surface soil moisture alone. Instead, they emerge from interactions among soil recharge, atmospheric energy demand, and stand composition. These interactions are missing in current Earth System Models, yet, they are essential for understanding the impacts of climate change on ecohydrology of mountain landscapes. Incorporating these processes will improve predictions of the future functioning of spruce forests in mountainous landscapes.

## Acknowledgements

HB acknowledges funding from the CryoSCOPE project (grant number: 101184736), supported by the State Secretariat for Education, Research and Innovation (SERI) under the European Union’s Horizon Europe research and innovation program. AS, MG and NB acknowledge funding from the ETH Zürich project FEVER (ETH-27 19-1), and the SNF funded projects ICOS-CH Phase 3 (20F120_198227) and EcoDrive (IZCOZ0_198094). We also thank Dillon Mungle for insightful discussions on mechanisms that could isotopically enrich deeper soil layers in the absence of growing-season soil recharge.

## Author contributions

NB and MG conceived the idea; HB, AS, NB, and MG designed the study; AS and MG conducted fieldwork and sampling; AS performed laboratory analysis; HB conducted the data analysis and wrote the first draft. All authors contributed substantially to reviewing, editing and revising the manuscript.

## Data availability

The raw isotope data supporting the findings of this study are provided in Supplementary Information. Upon acceptance, all data will be deposited in a publicly accessible repository. Figures S1-S5 and Notes S1 are available in the Supporting Information.

## Data Availability Statement

Our [Mandates Data Policy](https://authors.wiley.com/author-resources/Journal-Authors/open-access/data-sharing-citation/data-sharing-policy.html) requires data to be shared and a Data Availability Statement, so please enter one in the space below. Sample statements can be found [here](https://authors.wiley.com/author-resources/Journal-Authors/open-access/data-sharing-citation/data-sharing-policy.html#standardtemplates). Please note that this statement will be published alongside your manuscript, if it is accepted for publication.

The raw isotope data supporting the findings of this study are provided in Supplementary Information. Upon acceptance, all data will be deposited in a publicly accessibly repository. Figures S1-S5 and Notes S1 are available in the Supporting Information.

## Notes

### Competing Interest Statement

The authors have declared no competing interest.

